# Protective Effects of Lipoxin A_4_ and B_4_ Signaling on the Inner Retina in a Mouse Model of Experimental Glaucoma

**DOI:** 10.1101/2024.01.17.575414

**Authors:** Hsin-Hua Liu, Paul F. Cullen, Jeremy M. Sivak, Karsten Gronert, John G. Flanagan

**Affiliations:** Herbert Wertheim School of Optometry and Vision Science, University of California at Berkeley, Berkeley, California, United States; Department of Ophthalmology and Vision Science, University of Toronto, Toronto, Ontario, Canada; Krembil Research Institute, University Health Network, Toronto, Ontario, Canada

**Author notes:** **Corresponding Author:** John G. Flanagan, Herbert Wertheim School of Optometry and Vision Science, Minor Hall, University of California at Berkeley, Berkeley, CA 94720, USA, **Email:**. **Disclosures:** Jeremy M. Sivak, Karsten Gronert and John G. Flanagan are named on a patent titled “Lipoxin Mediated Neuroprotection and Treatments,” United States Patent Application No. 16/492,494.

## Abstract

Glaucoma is a common neurodegenerative disease characterized by progressive degeneration of retinal ganglion cells (RGCs) and the retinal nerve fiber layer (RNFL), resulting in a gradual decline of vision. A recent study by our groups indicated that the levels of lipoxins A_4_ (LXA_4_) and B_4_ (LXB_4_) in the retina and optic nerve decrease following acute injury, and that restoring their function is neuroprotective. Lipoxins are members of the specialized pro-resolving mediator (SPM) family and play key roles to mitigate and resolve chronic inflammation and tissue damage. Yet, knowledge about lipoxin neuroprotective activity remains limited. Here we investigate the *in vivo* efficacy of exogenous LXA_4_ and LXB_4_ administration on the inner retina in a mouse model of chronic experimental glaucoma. To investigate the contribution of LXA_4_ signaling we used transgenic knockout (KO) mice lacking the two mouse LXA_4_ receptors (Fpr2/Fpr3^-/-^). Functional and structural changes of inner retinal neurons were assessed longitudinally using electroretinogram (ERG) and optical coherence tomography (OCT). At the end of the experiment, retinal samples were harvested for immunohistological assessment. While both lipoxins generated protective trends, only LXB_4_ treatment was significant, and consistently more efficacious than LXA_4_ in all endpoints. Both lipoxins also appeared to dramatically reduce Müller glial reactivity following injury. In comparison, Fpr2/Fpr3 deletion significantly worsened inner retinal injury and function, consistent with an essential protective role for endogenous LXA_4_. Together, these results support further exploration of lipoxin signaling as a treatment for glaucomatous neurodegeneration.

## Introduction

Glaucoma is among the most common neurodegenerative diseases, and can lead to irreversible blindness. It is characterized by progressive degeneration of retinal ganglion cells (RGCs) and the retinal nerve fiber layer (RNFL), resulting in a gradual decline of visual function.^1, 2^ From a clinical perspective, the primary risk factors for glaucoma are aging and increased intraocular pressure (IOP). The standard of care for the disease is to lower IOP, which slows disease progression for many individuals.^3^ However, there is no treatment that directly targets RGC degeneration, and the mechanisms underlying this loss are not well understood.^4–6^ Glaucoma is associated with extensive glial reactivity and neuroinflammation, yet the role of glia in disease development are not fully understood.^7–10^ We and others have uncovered increasing evidence that astrocytes can also play essential roles in maintaining and promoting neuronal homeostasis.^11–16^

A recent study by our groups indicated that the levels of astrocyte-derived lipoxins A_4_ (LXA_4_) and B_4_ (LXB_4_) in the retina and optic nerve decreased following acute excitotoxic injury.^17^ Lipoxins are members of the specialized pro-resolving mediator (SPM) family of polyunsaturated fatty acid metabolites, derived from arachidonic acid.^18^ They play important roles as potent endogenous produced locally to mitigate and resolve chronic inflammation and tissue damage.^19–21^ Amplification of either LXA_4_ or LXB_4_ prior to excitotoxic retinal injury was able to promote RGC survival. In addition, *in vitro* results demonstrated that both LXA_4_ and LXB_4_ have direct protective effects on neuronal cells and RGCs in a dose-dependent fashion. Finally, we found that therapeutic administration of LXB_4_ was neuroprotective in a rat model of chronic experimental glaucoma, but LXA_4_ was not assessed at that time.

In a previous preclinical study, it had been reported that LXA_4_ could effectively dampen neuroinflammation in spinal cord injury by reducing pro-inflammatory cytokine release and suppressing microglial activation via the LXA_4_ receptor Fpr2 (formyl-peptide receptor 2). ^22, 23^ When compared to LXA_4_, which has been well documented, the role and function of LXB_4_ are much less well-studied and it’s receptor has not yet been identified. Nonetheless, LXB_4_ also displays the capacity to resolve spinal neuroinflammation in preclinical studies through a distinct pathway, independent of LXA_4_ signaling.^22, 24^ Yet, while evidence regarding the roles of lipoxins in neuroinflammation is robust, knowledge about their neuroprotective activity, especially in the retina and brain, remains relatively limited. As far as we know, our recent study was the first to demonstrate a direct link between lipoxins and neuroprotective bioactivity.^17^

In the present study, we investigated the *in vivo* effect of exogenous administration of LXA_4_ and role of the Fpr2/Fpr3^-/-^ receptors, in addition to LXB_4_ treatment, on the inner retina in a new mouse model of chronic experimental glaucoma. In order to clarify the contribution of canonical LXA_4_ signaling transgenic knockout (KO) mice lacking both LXA_4_ receptors, Fpr2/Fpr3^-/-^, and corresponding wild types were included in the study. Functional and structural changes of inner retinal neurons were assessed longitudinally using electroretinogram (ERG) and optical coherence tomography (OCT), respectively. In addition, retinal samples were also harvested for histological and immunohistological assessment.

## Materials and Methods

### Animals

All animals were treated in accordance with the ARVO Statement for the Use of Animals in Ophthalmic and Vision Research, and all experimental procedures were approved by the Animal Care and Use Committee of University of California at Berkeley. Six-week-old C57BL/6 mice (n=30 male, n=9 female) were purchased from the Jackson Laboratory (Bar Harbor, ME). The LXA_4_ receptor (Fpr2/Fpr3) KO mice on a C57BL/6 genetic background (n=8, 7∼8-week-old, female), were gifted by Dr. Asma Nusrat from the Michigan Medicine, University of Michigan (Ann Arbor, MI).^25^ All mice were allowed to acclimatize to the housing facility for two weeks prior to experimentation. Food and water were available ad libitum.

### Experimental Glaucoma

Experimental glaucoma was induced in all mice using a recently established rodent suture model of chronic ocular hypertension.^26–29^ Briefly, mice were anesthetized with ketamine/xylazine (100mg/kg and 10mg/kg, respectively, intraperitoneally). One drop of topical anesthetic (proparacaine hydrochloride, 0.5%, Akorn, Lake Forest, IL) was applied to either randomly selected eye. Body temperature of mice was maintained on a heating pad during procedure. To induce IOP elevation, a circumlimbal suture (10/0 nylon, Fine Science Tools, Foster City, CA) was secured subconjunctivally around the equator (0.4 ∼ 0.5 mm behind the limbus) on the selected eye by 5 anchor points and double knots on the conjunctiva. The tightness of the suture was controlled carefully to achieve desired IOP levels. Antibiotic ointment was applied to the eyes following the procedure. The contralateral untreated eye served as a within-animal control. A Tonolab rebound tonometer (iCare, Helsinki, Finland) was used to measure IOP by averaging 10 consecutive readings at baseline (1 day before surgery), immediately following surgery, and after 3 and 24 hours. After day 1, IOP was monitored weekly (between 10 am and 12 pm with identical lighting conditions) for 12 weeks. All readings were recorded in awake animals except for the reading immediately after surgery. The procedure was approximately 70% successful. Mice were excluded if their IOP peaked above 55mmHg immediately after suturing, or if IOP was below 25mmHg 1 day after suturing.

### Groupings and Treatments

Following ERG and OCT measurements at week 4, all male C57BL/6 mice were randomly treated with either vehicle (phosphate-buffered saline, PBS, n=10), LXA_4_ (n=10) or LXB_4_ (n=10). Both lipoxins were administered both systemically (5 ng/g, intraperitoneally; Cayman Chemical, Ann Arbor, MI) and topically to each eye (0.5 ng/g) every other day. New LXA_4_ and LXB_4_ PBS treatment solutions were prepared immediately before each treatment. The dosage and preparation of lipoxins was based on previous publications.^17, 30^ We chose to initiate therapeutic treatment following 4 weeks of IOP elevation since our previous work suggests this time point represents an early stage of retinal injury in the disease model.^27^ There might be increased potential for recovery of RGC dysfunction, as shown in a previous rat study.^31^ All Fpr2/Fpr3 KO mice (n=8) and sex & age-matched wild type (WT) C57BL/6 female mice (n=9) did not receive any therapeutic treatment throughout the experimental period.

### Electroretinography

Mice were dark-adapted overnight before ERG measurement. As described previously, they were anesthetized with ketamine & xylazine and topical proparacaine hydrochloride was used for corneal anesthesia. For pupil dilation, tropicamide (0.5%, Akorn, Lake Forest, IL) and phenylephrine (2.5%, Paragon BioTeck, Portland, OR) were applied. The VERIS system (Electro-Diagnostic Imaging, Redwood City, CA) was used for ERG measurement with a series of stimulus intensity ranging from -5.90 to 2.25 log cd.s.m^-2^. Silver corneal (positive) and scleral ring (reference) electrodes were custom-made as described previously.^26^ The tail subdermal needle electrode served as ground. The positive scotopic threshold response (pSTR) elicited with the intensity of -4.60 log cd.s.m^-2^ was used to represent RGC function (average of 20 repeats, inter-stimulus interval of 2 seconds). The amplitude at the time of 110 ms after stimulus onset was adopted.

### Optical Coherence Tomography

After ERG recording, mice were transferred to the OCT imaging platform. Lubricant eye drops (Systane Ultra, Alcon, Fort Worth, TX) were applied prior to imaging. A Bioptigen spectral domain OCT system (Envisu R2300, Durham, NC) was used. The OCT image acquisition and analysis protocols were described in our previous studies.^27, 29^ Briefly, an en-face retinal fundus image with the optic nerve head in the center was captured with a 3 x 3 mm rectangular scanning. Each image consisted of 100 B-scan images (1000 A-scans for each B-scan). Retinal layer B-scan images were analyzed with the InVivoVue Clinic software (caliper function) by masked observers. Retinal layer thickness was quantified at 3 locations (400, 500 and 600 μm from the center of the optic nerve head) in each retinal quadrant. The average of data from all 4 retinal quadrants was used to represent the thickness value.

### Retinal Flat Mount and RGC Quantification

For ocular tissue collection, mice were euthanized by carbon dioxide inhalation and cervical dislocation at week 12 following OCT imaging. The tissue process procedure and analysis protocol were performed as described previously.^27, 29^ In brief, eyes were enucleated, fixed with 4% paraformaldehyde (room temperature, 15 minutes) and washed in PBS (10 minutes). The eyeball was bisected by an incision on the edge of cornea. The cup-shaped retina was isolated and then flattened by 4 radial cuts while rinsed in PBS. Following fixation with methanol (-20°C, overnight), the retina was rinsed with PBS (30 minutes for 3 times at room temperature). RGCs were labeled by incubation with a primary goat antibody raised against Brn3a (brain-specific homeobox/POU domain protein 3A, Santa Cruz Biotechnology, Santa Cruz, CA), 1:100 in PBS with 2% bovine serum albumin, 2% Triton, at 4°C overnight with gentle shaking. Alexa Fluor donkey anti-goat IgG (1:200, room temperature, 2 hours; Jackson ImmunoResearch Laboratories, West Grove, PA) was used as a secondary antibody. Retinal samples were mounted on slides with an antifading medium (Prolong Gold, Invitrogen, Carlsbad, CA). A Zeiss Axioplan epifluorescent microscope system (Carl Zeiss, Oberkochen, Germany) was used to capture retinal images for RGC quantification. A total of 8 areas (each 450 x 320 µm in size, 20x, 850 µm from the center of the optic nerve head) from all retinal quadrants were imaged. The RGC density in each area was evaluated with ImageJ software (National Institutes of Health, Bethesda, MD) by masked observers. RGC density data from all areas were averaged to return a value for that retina.

### Immunofluorescence Staining for Glial Reactivity

After enucleation, the anterior segment and lens were removed and eyes were fixed in 4% paraformaldehyde (PFA) for 15 minutes at room temperature. Fixative was removed by wash with PBS, and eyes were subsequently equilibrated in 30% sucrose before flash-freezing to embed in optimal temperature compound. Eyes were cryosectioned and sections rinsed in PBS before blocking and permeabilization for 1 hour at room temperature in PBS with 0.1% Triton X-100 and 10% goat serum. Sections were then stained overnight at room temperature with 1:1000 rabbit anti-GFAP primary antibody (Abcam, AB7260), and after additional PBS rinses, stained for 1 hour with 1:300 goat-anti-rabbit secondary (Invitrogen, A-11008), in PBS with 0.1% Triton X-100 and 3% goat serum. After an additional PBS rinse, nuclear counterstaining was performed with 1 µg/ml Hoechst stain for 20 minutes before a final PBS wash. Images were captured with a Nikon Eclipse-Ti confocal microscope (Nikon Inc., Melville, NY).

### Statistical Analyses

All data were expressed as mean ± SEM. Prism 6 software (GraphPad, La Jolla, CA) was used for data analysis and statistics. IOP, ERG and OCT data were analyzed using a repeated measures (RM) two-way ANOVA with Bonferroni post hoc test. For RGC loss data, a one-way ANOVA with Bonferroni multiple comparisons test or unpaired t-test was used, where applicable. The percentage loss was calculated as: ((treated – control) / control) x 100%.

## Results

The IOP data following model induction and treatment are shown in Figure 1 (n = 10 for each group). In all groups, IOP was significantly elevated in sutured eyes when compared to that in control eyes. In all sutured eyes, a two-way RM ANOVA showed that there was no significant interaction between group and time (p = 0.62) and no significant group effect (p = 0.15). A similar result was noted for all control eyes. These results indicate that all 3 groups had a similar IOP profile over time.

**Figure 1.**
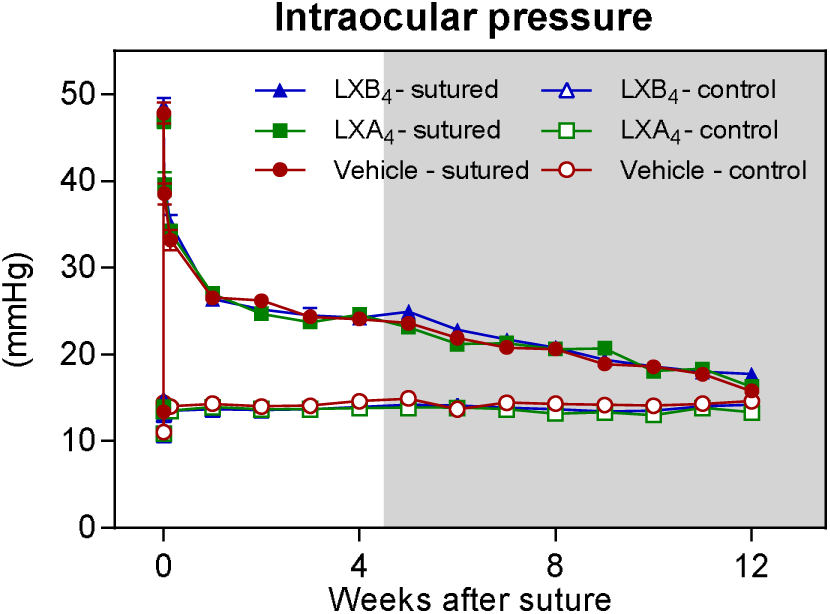
IOP was similarly elevated under each treatment condition. Significant ocular hypertension was noted in all sutured eyes and no significant change in IOP was found between the three groups across 12 weeks. The shaded area indicates the period of treatment. (Markers are mean ± SEM, n = 10 for each group).

Changes in RGC function were measured longitudinally by ERG measurements of pSTR (n = 10 for each group) (Figure 2). The average waveforms of pSTR in were elicited by a flash intensity of -4.60 log cd.s.m^-2^ at 12 weeks (Figure 2A). A relative loss of RGC function appears to progressive over 12 weeks in all groups (Figure 2B). There was a significant statistical interaction between group and time (p < 0.01). In the vehicle group the loss was - 31.1 ± 1.8% at week 12. For the LXA_4_ treatment group, the loss was -26.1 ± 1.3% and for the LXB_4_ group the loss was -23.9 ± 1.4% (Figure 2B). Although both lipoxin treatments reduced ocular hypertension-indued RGC loss over time, a significant difference was noted only between the vehicle group and LXB_4_ group (p < 0.01).

**Figure 2.**
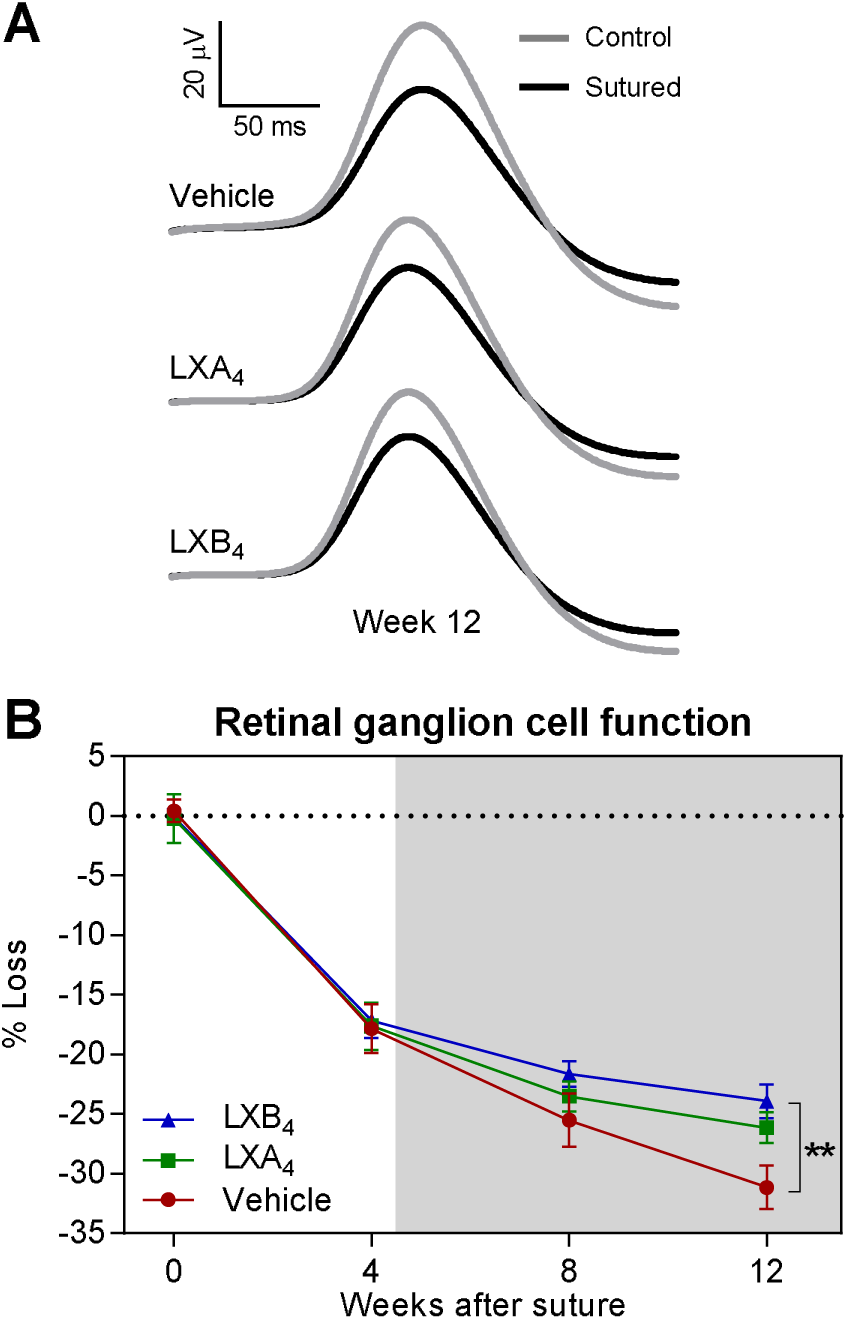
Therapeutic lipoxin treatment reduces loss of RGC function. (A) Average ERG waveforms of RGC responses (pSTR, -4.60 log cd.s.m^-2^) at week 12 for each group. (B) A progressive loss of RGC function was found in all groups; however, the loss in the LXB_4_ group was significantly alleviated (**p < 0.01) at week 12 when compared to the vehicle group. The shaded area indicates the period of treatment. (Markers are mean ± SEM, n = 10 for each group).

In parallel to functional measures, longitudinal alterations in RNFL thickness were measured by OCT, as a clinically relevant biomarker of nerve fiber loss (Figure 3). The RNFL thickness of sutured eyes in the vehicle group at week 12 was 15.8 ± 0.9 μm. In the LXA_4_ group, it was 16.2 ± 0.9 μm, and in the LXB_4_ group it was 17.2 ± 0.7 μm (Figure 3A). When compared to contralateral control eyes, the relative loss was -25.2 ± 1.4% in the vehicle group, -21.8 ± 1.6% in the LXA_4_ group and -18.4 ± 1.5% in the LXB_4_ group (Figure 3B). Similar to the ERG measures both treatments reduced RNFL loss, but a significant difference was noted only between the vehicle group and LXB_4_ group (p < 0.01).

**Figure 3.**
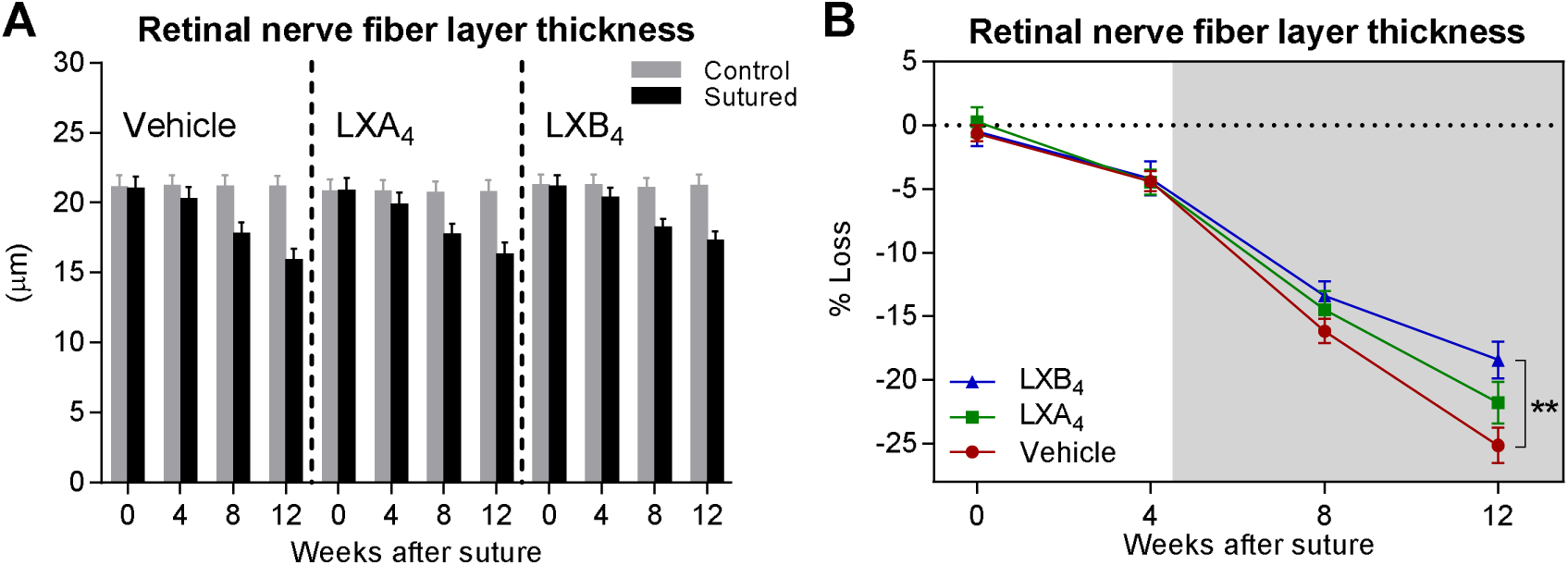
RNFL thickness is preserved by lipoxin treatment. (A) Structural changes in the RNFL were monitored using OCT across 12 weeks. (B) The loss of RNFL was progressive for all groups. At week 12, the loss in the LXB_4_ group was significantly mitigated (** p < 0.01) when compared to that in the vehicle group. The shaded area indicates the period of treatment. (Markers are mean ± SEM, n = 10 for each group).

Finally, after twelve weeks all animals were euthanized and nine of the ten retinas were collected for flatmounting and counting of RGC density (Figure 4A). At week 12, the relative RGC loss for the vehicle group was -17.5 ± 1.3%, whereas it was -14.0 ± 1.1% for the LXA_4_ treatment group (p = 0.13) and -12.2 ± 1.0% for the LXB_4_ treatment group (p < 0.01; Figure 4B).

**Figure 4.**
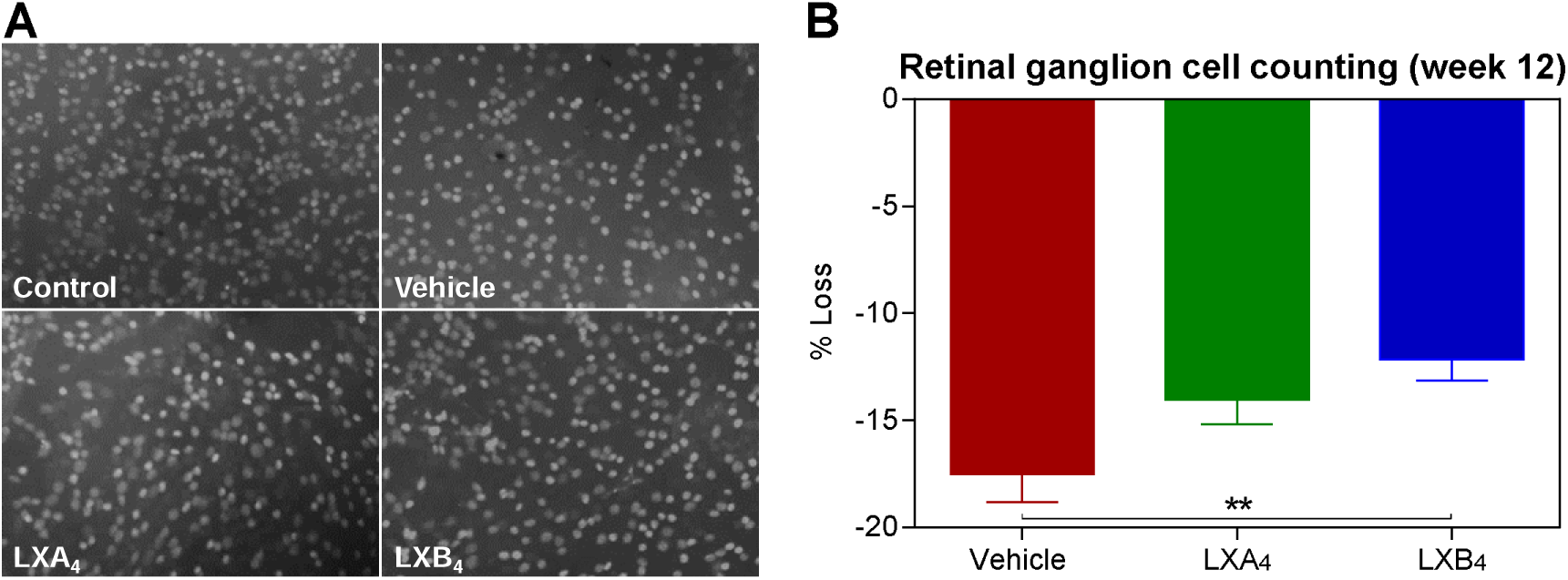
RGC loss is reduced by lipoxin treatment. (A) Representative flatmount Brn3a-labeled RGC images (450 x 320 μm) at week 12. (B) When compared to the vehicle group, the LXB_4_ treatment group displayed significantly less RGC loss (** p < 0.01). (Bars are mean ± SEM, n = 9 for each group).

We have identified retinal ganglion cells as a primary target for the protective actions of endogenous lipoxins and glia reactivity as an important regulator of the lipoxin pathway in the retina.^17^ In order to assess the effect of our model on glial reactivity, and how lipoxin treatment may modify reactivity, the eyes from the remaining animal in each group of ten were frozen and sectioned for immunofluorescent staining. Staining for the intermediate filament astrocyte marker GFAP (glial fibrillary acidic protein) showed similar localization in the RNFL across control and all treatment conditions. However, retinal sections from the vehicle treated eye demonstrated that ocular hypertension induced marked expression of GFAP in Müller glia as well, which is a marker of glia reactivity. Müller glia in unsutured control eyes did not express GFAP. More importantly, GFAP expression was absent or markedly reduced in LXA_4_ or LXB_4_ treated eyes (Figure 5), consistent with reduced Müller glia reactivity.

**Figure 5.**
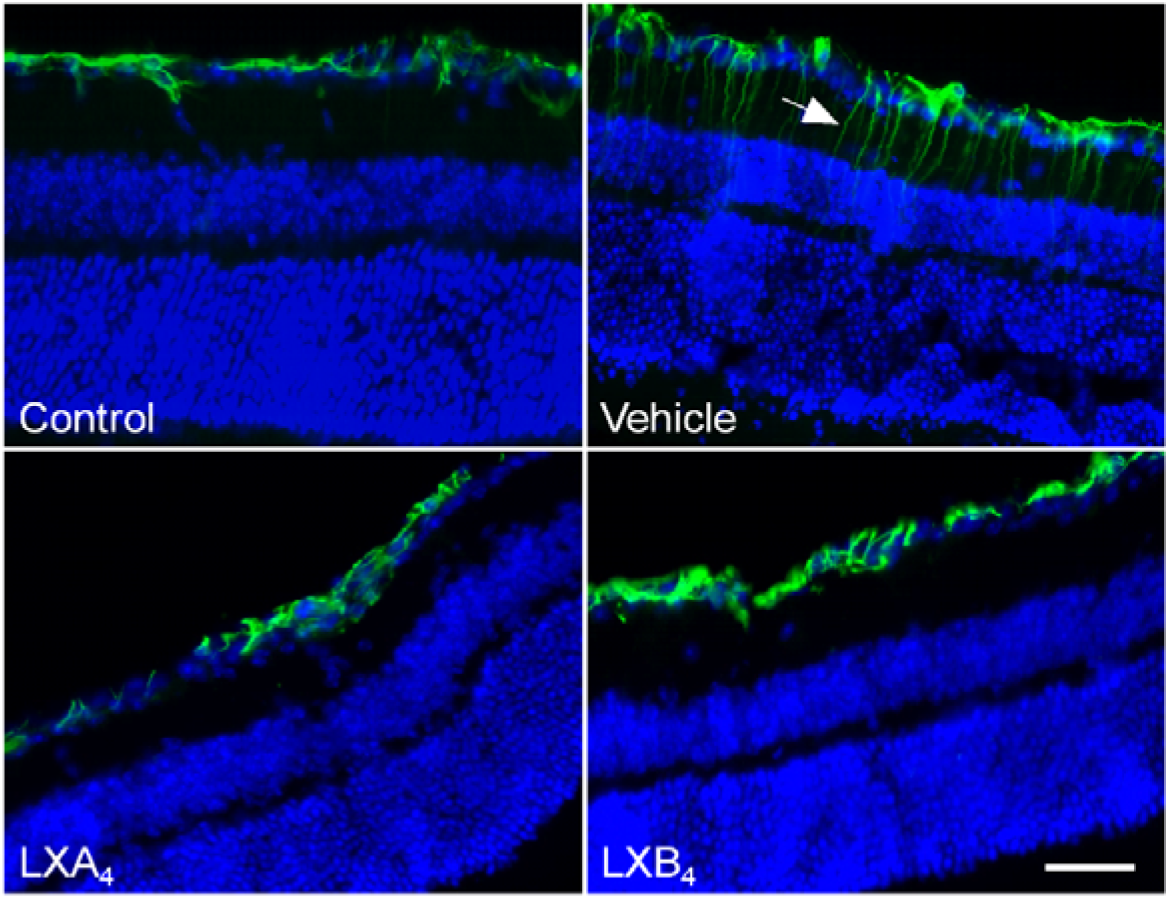
Induced Müller glia reactivity was inhibited by lipoxin treatment. At week 12, eyes of one animal from each group were cryosectioned and immunostained with anti-GFAP (green). Induced GFAP expression in Müller cell processes in the inner plexiform layer were abundantly noted in the vehicle treated eye (arrow), but were not found in the unsutured control, LXA_4_ or LXB_4_ treated eyes . Scale bar = 50 µm.

Similar to our previously published *in vitro* results,^17^ the present experiments indicated a stronger significant protective effect for LXB_4_ treatment compared to LXA_4_, which often did not reach significance under our current therapeutic dose and regiment. Therefore, as an alternate strategy to study the role of endogenous LXA_4_ signaling in our mouse glaucoma model, we assessed IOP and RGC parameters in mice carrying a genetic deletion of both mouse LXA_4_ receptors, namely Fpr2 and Fpr3.^25^ Notably, the IOP profile of the sutured eyes from the WT group (n = 9) and the Fpr2/Fpr3^-/-^ group (n = 8) was similar, indicating no effect of Fpr2 or Fpr3 signaling on baseline IOP or the suture induced IOP response (Figure 6A; interaction p = 0.36, group p = 0.37). RGC function was measured by ERG, with a progressive loss of function observed across 12 weeks for both groups. However, the loss at week 12 was -36.7 ± 1.6% in the WT group and increased to -45.4 ± 2.5% in the Fpr2/Fpr3^-/-^ group (p < 0.01) (Figure 6B). A corresponding reduced RNFL thickness was also observed by OCT measurement for knockout eyes (Figure 6C). Reduction in RNFL thickness at week 12 was -29.2 ± 2.0% for the WT group and -40.0 ± 2.8% for LXA_4_ receptor deficient (Fpr2/Fpr3^-/-^) group (p < 0.01). Finally, the loss of RGC density was -25.3 ± 1.6% in the WT group and increased to -34.8 ± 2.3% in Fpr2/Fpr3^-/-^ eyes (p < 0.01, unpaired t-test; Figure 6D). Together, these observations indicate that Fpr2/Fpr3 signaling has a protective role for inner retinal responses to chronic increased IOP injury.

**Figure 6.**
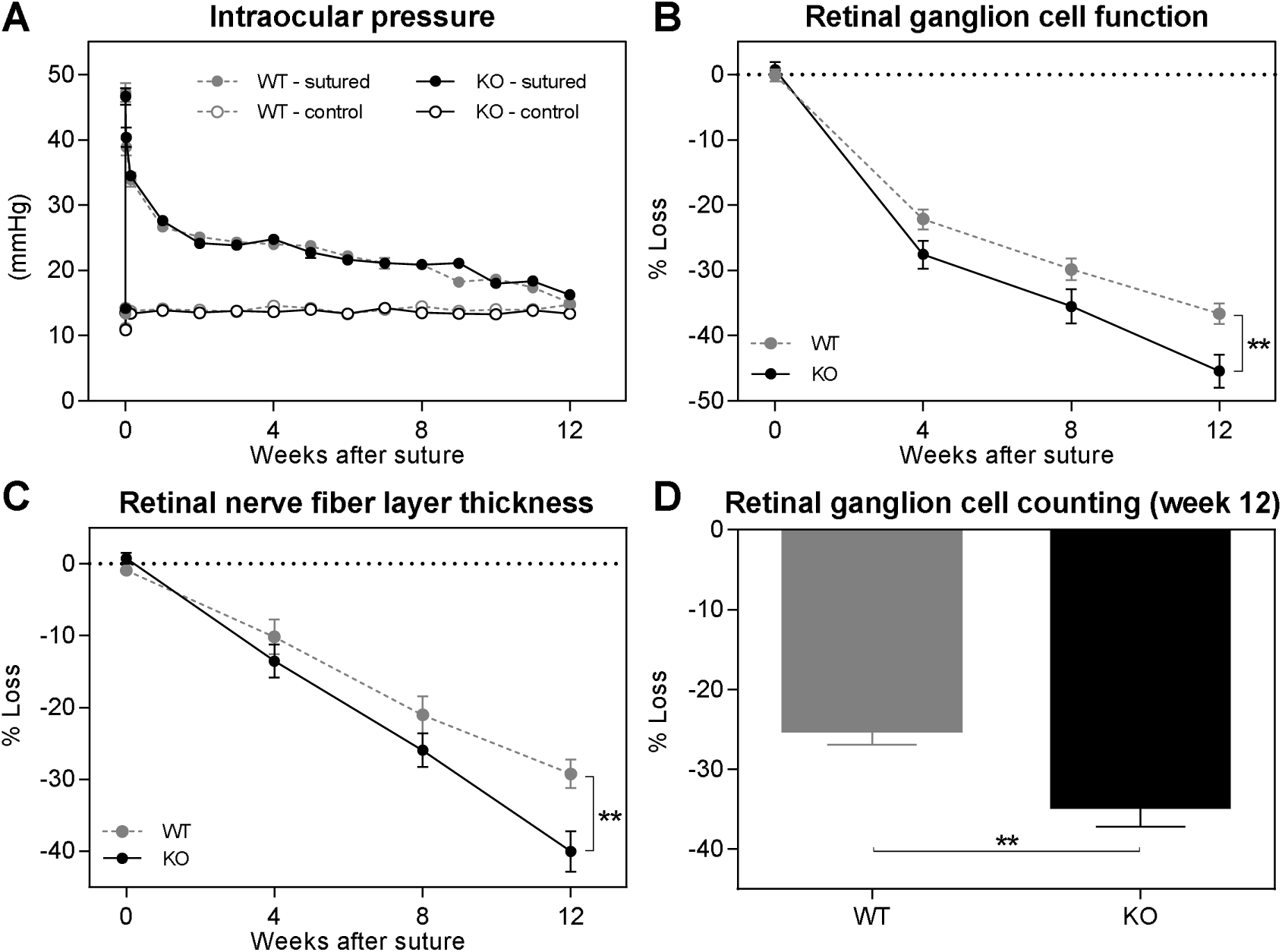
ALX/FPR2 is essential for normal inner retinal glaucomatous injury responses. (A) A similar IOP profile was found between the WT and ALX/FPR2^-/-^ eyes. At week 12, the loss of RGC function (B), RNFL thickness (C) and RGC density (D) were significantly greater in the ALX/FPR2^-/-^ group compared to those in the WT group. (Markers and bars are mean ± SEM, ** p < 0.01, n = 9 in the WT group, n = 8 in the KO group).

## Discussion

Our data indicate that therapeutic administration of LXB_4_ provides marked neuroprotection in mouse experimental glaucoma determined by clinical relevant measures, via a mechanism independent of IOP. LXA_4_ treatment consistently showed similar trends but did not reach statistical significance. However, deletion of both LXA_4_ receptors, Fpr2 and Fpr3, resulted in significantly worse inner retinal injury in experimental glaucoma, providing strong evidence for a protective role for the LXA_4_ signaling pathway. Furthermore, the resulting lipoxin-induced neuroprotective mechanism may be associated with inhibiting glial cell reactivity in the retina.

Glaucoma is a complex neurodegenerative disease with a multifactorial origin.^5^ In addition to primary contributing factors such as high IOP, secondary pathological pathways including glial dysregulation and neuroinflammation driven by glial immunoreactivity have been shown to play a role in the pathogenesis of the disease.^32–35^ Given glaucoma may still advance following IOP-lowering treatments, investigations targeting the molecular mechanisms associated with neurodegeneration have emerged for the development of neuroprotection strategies.^36, 37^ In the present study, we found that exogenous administration of the LXB_4_, best known for its anti-inflammatory and pro-resolving activities, mitigated the functional and structural loss in experimental glaucoma by a non-IOP-lowering mechanism. This finding is consistent with our previous rat glaucoma study.^17^ Moreover, in our mouse model LXB_4_ appeared to consistently exhibit greater efficacy than LXA_4_ in all of our outcome measures (Figures 2-4), which is also in agreement with our previous in vitro and in vivo experiments.^17^ These result are of particular interest since LXB_4_ bioactivity, formation and receptor identity are not well understood in comparison with LXA_4_. Therefore, it is essential to uncover the LXB_4_ signaling pathway for the development of its potential therapeutic applications.

Growing evidence has suggested an immunomodulation strategy promoting neuroprotection in glaucoma via resolution of neuroinflammation.^22, 38–41^ It is of interest that lipoxins promote resolution of neuroinflammation by inhibition of pro-inflammatory cytokine release^23, 24^. More importantly, our previous study^17^ identified direct actions of LXB_4_ with RGC and primary neuron as a novel neuroprotective mechanism of action for lipoxins. Hence, lipoxins and their signaling pathways are well positioned as promising therapeutic targets for the treatment of glaucoma. A receptor for LXB_4_ has not been identified but our data shows that LXA_4_ receptor deficiency (Fpr2/Fpr3^-/-^) leads to enhanced RGC dysfunction and death (Figure 6). Treatment with LXA_4_, unlike LXB_4_, in the present study produced a positive trend, but did not achieve statistically significant neuroprotection. However, data from the Fpr2/Fpr3^-/-^ mice provides strong evidence for a neuroprotective role of the resident retinal LXA_4_ receptors.^17^ A limitation for LXA_4_ treatment efficacy maybe the relatively late therapeutic intervention in our model of experimental glaucoma. At week 4, there was already ∼18% loss of RGC function in all groups, accounting for much of the total RGC functional loss at the end of the experimental period (12 weeks) in the vehicle group.

As a result, therapeutic treatment with LXA_4_ from week 5 likely resulted in diminished statistical power to assess functional rescue. LXB_4_ does not act at the LXA_4_ receptors and each lipoxin has distinct signaling pathways in leukocyte.^42^ It is likely that LXA_4_ and LXB_4_ have distinct and temporally defined protective action in the retina and pathogenesis of neurodegeneration, which may explain the difference between the efficacy of LXB_4_ and LXA_4_ treatment and the endogenous role of LXA_4_ receptor signaling. Our previous *in vitro* studies demonstrated that both lipoxins have direct neuroprotective activities with oxidative stress and glutamate-challenged neuronal cells, including RGCs.^17^ Hence, based on the endogenous protective role of LXA_4_ receptor signaling and direct action with RGCs, we hypothesize that an earlier treatment with LXA_4_ or LXA_4_ stable mimetics might lead to improved therapeutic outcomes similar to those of LXB_4_.

In addition to the functional and pathological RGC outcomes in our study, we observed that both LXA_4_ and LXB_4_ treatment inhibited the induction of GFAP in Müller glia fibers following experimental glaucoma (Figure 5). Unlike retinal astrocytes, which constitutively express GFAP and are generally restricted to the RNFL, Müller glia traverse the retina and typically do not exhibit GFAP expression outside of disease or injury states.^43^ A previous characterization of our disease model showed increased GFAP expression in reactive Müller glia, leading us to investigate the effect lipoxin treatment might have in the present study.^26^ As expected, Müller glia in vehicle treated eyes demonstrated GFAP expression while those in the contralateral normotensive control eye did not. Intriguingly, the GFAP induction was absent in both LXA_4_ and LXB_4_ treated eyes, suggesting that in addition to providing neuroprotective effects, these lipoxins also reduce Müller glia reactivity, either directly or as a result of improved RGC survival. Further investigation is required to elucidate the relationship between these changes.

In summary, we have identified distinct and new neuroprotective actions for endogenous lipoxins, LXA_4_ and LXB_4_, and the LXA_4_ receptors (Fpr2 and Fpr3) in a new chronic ocular hypertension mouse glaucoma model. Therapeutic treatment with lipoxins markedly reduced RGC loss and prevented Müller glia cell reactivity. Genetic deletion of the two LXA_4_ receptors identified a necessary role for LXA_4_ signaling in limiting glaucoma pathogenesis. The current findings provide strong evidence that lipoxin signaling presents potential therapeutic targets for the treatment of glaucomatous neurodegeneration.

## Acknowledgements

Supported by NIH grant R01EY030218 (KG, JGF, and JMS), and a Shaffer Grant from the Glaucoma Research Foundation (JGF). JMS holds the TGWHF Glaucoma Research Chair.

## References

[1] Kwon YH, Fingert JH, Kuehn MH, Alward WL: Primary open-angle glaucoma. The New England journal of medicine 2009, 360:1113–24.

[2] Weinreb RN, Khaw PT: Primary open-angle glaucoma. Lancet 2004, 363:1711–20.

[3] Weinreb RN, Aung T, Medeiros FA: The pathophysiology and treatment of glaucoma: a review. Jama 2014, 311:1901–11.

[4] Almasieh M, Wilson AM, Morquette B, Cueva Vargas JL, Di Polo A: The molecular basis of retinal ganglion cell death in glaucoma. Prog Retin Eye Res 2012, 31:152–81.

[5] Alqawlaq S, Flanagan JG, Sivak JM: All roads lead to glaucoma: Induced retinal injury cascades contribute to a common neurodegenerative outcome. Exp Eye Res 2019, 183:88–97.

[6] Tezel G: A proteomics view of the molecular mechanisms and biomarkers of glaucomatous neurodegeneration. Prog Retin Eye Res 2013, 35:18–43.

[7] Johnson EC, Morrison JC: Friend or foe? Resolving the impact of glial responses in glaucoma. J Glaucoma 2009, 18:341–53.

[8] Mac Nair CE, Nickells RW: Neuroinflammation in Glaucoma and Optic Nerve Damage. Prog Mol Biol Transl Sci 2015, 134:343–63.

[9] Russo R, Varano GP, Adornetto A, Nucci C, Corasaniti MT, Bagetta G, Morrone LA: Retinal ganglion cell death in glaucoma: Exploring the role of neuroinflammation. Eur J Pharmacol 2016, 787:134–42.

[10] Williams PA, Marsh-Armstrong N, Howell GR, Lasker IIoA, Glaucomatous Neurodegeneration P: Neuroinflammation in glaucoma: A new opportunity. Exp Eye Res 2017, 157:20–7.

[11] Alqawlaq S, Livne-Bar I, Williams D, D’Ercole J, Leung SW, Chan D, Tuccitto A, Datti A, Wrana JL, Corbett AH, Schmitt-Ulms G, Sivak JM: An endogenous PI3K interactome promoting astrocyte-mediated neuroprotection identifies a novel association with RNA-binding protein ZC3H14. J Biol Chem 2021, 296:100118.

[12] Guo X, Jiang Q, Tuccitto A, Chan D, Alqawlaq S, Won GJ, Sivak JM: The AMPK-PGC-1α signaling axis regulates the astrocyte glutathione system to protect against oxidative and metabolic injury. Neurobiol Dis 2018, 113:59–69.

[13] Nahirnyj A, Livne-Bar I, Guo X, Sivak JM: ROS detoxification and proinflammatory cytokines are linked by p38 MAPK signaling in a model of mature astrocyte activation. PLoS One 2013, 8:e83049.

[14] Sofroniew MV: Astrogliosis. Cold Spring Harb Perspect Biol 2015, 7:a020420.

[15] Sofroniew MV, Vinters HV: Astrocytes: biology and pathology. Acta Neuropathol 2010, 119:7–35.

[16] Sun D, Moore S, Jakobs TC: Optic nerve astrocyte reactivity protects function in experimental glaucoma and other nerve injuries. J Exp Med 2017, 214:1411–30.

[17] Livne-Bar I, Wei J, Liu HH, Alqawlaq S, Won GJ, Tuccitto A, Gronert K, Flanagan JG, Sivak JM: Astrocyte-derived lipoxins A4 and B4 promote neuroprotection from acute and chronic injury. The Journal of Clinical Investigation 2017, 127:4403–14.

[18] Serhan CN, Hamberg M, Samuelsson B: Lipoxins: novel series of biologically active compounds formed from arachidonic acid in human leukocytes. Proc Natl Acad Sci U S A 1984, 81:5335–9.

[19] Serhan CN: Resolution phase of inflammation: novel endogenous anti-inflammatory and proresolving lipid mediators and pathways. Annu Rev Immunol 2007, 25:101–37.

[20] Ryan A, Godson C: Lipoxins: regulators of resolution. Curr Opin Pharmacol 2010, 10:166–72.

[21] Chandrasekharan JA, Sharma-Walia N: Lipoxins: nature’s way to resolve inflammation. J Inflamm Res 2015, 8:181–92.

[22] Kim C, Livne-Bar I, Gronert K, Sivak JM: Fair-Weather Friends: Evidence of Lipoxin Dysregulation in Neurodegeneration. Mol Nutr Food Res 2020, 64:e1801076.

[23] Martini AC, Berta T, Forner S, Chen G, Bento AF, Ji RR, Rae GA: Lipoxin A4 inhibits microglial activation and reduces neuroinflammation and neuropathic pain after spinal cord hemisection. J Neuroinflammation 2016, 13:75.

[24] Hu S, Mao-Ying QL, Wang J, Wang ZF, Mi WL, Wang XW, Jiang JW, Huang YL, Wu GC, Wang YQ: Lipoxins and aspirin-triggered lipoxin alleviate bone cancer pain in association with suppressing expression of spinal proinflammatory cytokines. J Neuroinflammation 2012, 9:278.

[25] Dufton N, Hannon R, Brancaleone V, Dalli J, Patel HB, Gray M, D’Acquisto F, Buckingham JC, Perretti M, Flower RJ: Anti-inflammatory role of the murine formyl-peptide receptor 2: ligand-specific effects on leukocyte responses and experimental inflammation. J Immunol 2010, 184:2611–9.

[26] Liu HH, Bui BV, Nguyen CT, Kezic JM, Vingrys AJ, He Z: Chronic ocular hypertension induced by circumlimbal suture in rats. Investigative ophthalmology & visual science 2015, 56:2811–20.

[27] Liu HH, Flanagan JG: A Mouse Model of Chronic Ocular Hypertension Induced by Circumlimbal Suture. Investigative ophthalmology & visual science 2017, 58:353–61.

[28] Zhao D, Nguyen CT, Wong VH, Lim JK, He Z, Jobling AI, Fletcher EL, Chinnery HR, Vingrys AJ, Bui BV: Characterization of the Circumlimbal Suture Model of Chronic IOP Elevation in Mice and Assessment of Changes in Gene Expression of Stretch Sensitive Channels. Frontiers in neuroscience 2017, 11:41.

[29] Liu HH, Zhang L, Shi M, Chen L, Flanagan JG: Comparison of laser and circumlimbal suture induced elevation of intraocular pressure in albino CD-1 mice. PLoS One 2017, 12:e0189094.

[30] Borgeson E, Johnson AM, Lee YS, Till A, Syed GH, Ali-Shah ST, Guiry PJ, Dalli J, Colas RA, Serhan CN, Sharma K, Godson C: Lipoxin A4 Attenuates Obesity-Induced Adipose Inflammation and Associated Liver and Kidney Disease. Cell Metab 2015, 22:125–37.

[31] Liu HH, He Z, Nguyen CT, Vingrys AJ, Bui BV: Reversal of functional loss in a rat model of chronic intraocular pressure elevation. Ophthalmic Physiol Opt 2017, 37:71–81.

[32] Krizaj D, Ryskamp DA, Tian N, Tezel G, Mitchell CH, Slepak VZ, Shestopalov VI: From mechanosensitivity to inflammatory responses: new players in the pathology of glaucoma. Curr Eye Res 2014, 39:105–19.

[33] Nickells RW, Howell GR, Soto I, John SW: Under pressure: cellular and molecular responses during glaucoma, a common neurodegeneration with axonopathy. Annu Rev Neurosci 2012, 35:153–79.

[34] Tang J, Tang Y, Yi I, Chen DF: The role of commensal microflora-induced T cell responses in glaucoma neurodegeneration. Prog Brain Res 2020, 256:79–97.

[35] Tezel G: Immune regulation toward immunomodulation for neuroprotection in glaucoma. Curr Opin Pharmacol 2013, 13:23–31.

[36] He S, Stankowska DL, Ellis DZ, Krishnamoorthy RR, Yorio T: Targets of Neuroprotection in Glaucoma. J Ocul Pharmacol Ther 2018, 34:85–106.

[37] Levin LA, Crowe ME, Quigley HA, Lasker IIoA, Glaucomatous Neurodegeneration P: Neuroprotection for glaucoma: Requirements for clinical translation. Exp Eye Res 2017, 157:34–7.

[38] Cueva Vargas JL, Belforte N, Di Polo A: The glial cell modulator ibudilast attenuates neuroinflammation and enhances retinal ganglion cell viability in glaucoma through protein kinase A signaling. Neurobiol Dis 2016, 93:156–71.

[39] Lambert WS, Carlson BJ, Formichella CR, Sappington RM, Ahlem C, Calkins DJ: Oral Delivery of a Synthetic Sterol Reduces Axonopathy and Inflammation in a Rodent Model of Glaucoma. Frontiers in neuroscience 2017, 11:45.

[40] Madeira MH, Ortin-Martinez A, Nadal-Nicolas F, Ambrosio AF, Vidal-Sanz M, Agudo-Barriuso M, Santiago AR: Caffeine administration prevents retinal neuroinflammation and loss of retinal ganglion cells in an animal model of glaucoma. Sci Rep 2016, 6:27532.

[41] Yang X, Hondur G, Tezel G: Antioxidant Treatment Limits Neuroinflammation in Experimental Glaucoma. Investigative ophthalmology & visual science 2016, 57:2344–54.

[42] Romano M, Maddox JF, Serhan CN: Activation of human monocytes and the acute monocytic leukemia cell line (THP-1) by lipoxins involves unique signaling pathways for lipoxin A4 versus lipoxin B4: evidence for differential Ca2+ mobilization. J Immunol 1996, 157:2149–54.

[43] Chang ML, Wu CH, Jiang-Shieh YF, Shieh JY, Wen CY: Reactive changes of retinal astrocytes and Muller glial cells in kainate-induced neuroexcitotoxicity. J Anat 2007, 210:54–65.

